# Visual Salience Controls the Speed of Evidence Accumulation in Value-Based Decisions by Rats

**DOI:** 10.1101/2025.10.24.684442

**Authors:** Jensen A Palmer, Kevin Chavez Lopez, Mark Laubach

## Abstract

Studies of visual discrimination in rodents can confound the effects of cue salience with reward value, making it difficult to determine which factor guides choice behavior. We examined this issue by testing how changes in relative salience affect decision dynamics in rats performing a two-alternative forced-choice task in which rats chose between visual cues associated with high or low sucrose rewards. After initial training with high and low luminance cues, we introduced a novel cue of intermediate luminance as a “luminance shift” test. The intermediate luminance cue substituted for either the brighter or dimmer cue and had the same reward value as the cue that it replaced. We found that while rats maintained a preference for the higher-value option, the introduction of a perceptually more similar cue consistently reduced choice preference and eliminated latency differences compared to baseline. Using drift diffusion modeling, we determined that the luminance shifts primarily caused a reduction in the drift rate (the speed of evidence accumulation), reflecting increased difficulty in cue discrimination. This finding suggests that the relative salience of the options determines the efficiency of evidence accumulation in value-based decisions. Furthermore, this effect on drift rate shows a dissociation from our previous work (Palmer et al., 2024), where prefrontal cortex inactivation specifically affected only the decision threshold. Our results demonstrate that relative salience influences deliberation, with low-level perceptual features shaping the computational dynamics of value-based choice. Our findings clarify the distinct contributions of sensory input and prefrontal function in the decision process.

**Significance Statement:** This study reveals that changes in the relative salience of visual stimuli shape the computational dynamics of value-based decisions. We trained rats to make visually guided choices and found that relative differences in the brightness of the stimuli affect how quickly the rats made decisions and how often they chose a higher-value option. Our findings, together with a recent study on the role of the prefrontal cortex in value-guided decisions (Palmer et al., 2024), suggest that separate factors influence choice dynamics in rodents: visual salience affects the speed of deliberation, while prefrontal activity regulates caution. This study helps clarify how sensory and higher cognitive variables relate to the distinct computational components of the decision process.

## Introduction

The specific design of a behavioral task has a significant impact on how we interpret its results. One key factor contributing to variations across studies from different research groups is the sensory modality used, such as visual (Zoccolan et al., 2009), auditory (Brunton et al., 2013), and olfactory (Vazquez et al., 2024) cues. As the time scales for processing sensory information differ across modalities, it is challenging to compare findings between studies. Visual stimuli are commonly used in decision-making research involving humans (e.g., Tanner and Swets, 1954) and non-human primates (e.g., Newsome et al., 1989). However, rodents have been understudied for their ability to perform visually guided behavioral tasks. Early studies (e.g., Lashley, 1912) demonstrated that rats could perform tasks based on the form of visual objects. Although research continued on rodents’ visual capabilities (e.g., Fields, 1928; Gentry, 1934; Wolfle, 1937), most studies on visual processing used simple stimuli, such as illuminating incandescent lights (e.g., Rescorla, 1990). These studies primarily focused on how rodents learned tasks by responding to simple cues, rather than exploring how the properties of the cues influenced their responses.

Rodents have become popular subjects for studying perceptual function and decision-making, driven by advances like optogenetics (Carandini & Churchland, 2013). A renewed interest in understanding rodent vision led to several key findings. For example, a study by Zoccolan et al. (2009) demonstrated that rats are capable of higher-level visual functions, specifically invariant object recognition. Subsequent studies have demonstrated that rodents are capable of visual discrimination based on contrast sensitivity (Busse et al., 2011), the shape of images (Clark et al., 2011), local patterns of luminance and texture (De Keyser et al., 2015), dynamically fluctuating patterns of luminance (Swanson et al., 2021), and local features in images (Schnell et al., 2023). These studies have established some of the basic properties of rodent vision for perceptual decisions, but have not explored how the properties of visual stimuli contribute to value-guided decisions.

Our group designed a two-alternative forced-choice (2AFC) task to investigate visually guided decisions (White et al., 2024; Palmer et al., 2024). The task used dynamic LED stimuli (Swanson et al., 2021) where brighter cues (16 LEDs) yielded larger rewards (16% sucrose), and dimmer cues (1 LED) yielded smaller rewards (1% sucrose). Rats responded faster to and chose the brighter, higher-value cue more often (∼80% preference). A potential interpretational problem with this design is that the rats could have responded to cues because they were brighter, and thus more salient, or because they had a higher reward value.

The present study addresses this issue by using “luminance shift” tests to examine how changes in relative salience affect decision dynamics within a fixed reward structure. Inspired by the design of a study of Go/No-Go performance (Kimchi and Laubach, 2009), animals were trained on a two-alternative forced choice task to choose between a high-luminance cue (16 LEDs/16% sucrose) and a low-luminance cue (1 LED/1% sucrose). After training, we introduced an intermediate luminance cue (4 LEDs) for the luminance shift phase. In shifting blocks, the animals were tested on a new pair: either the Upshift (High and Low→High and Intermediate) or the Downshift (High and Low→Intermediate and Low). Importantly, the reward values remained constant throughout the shift phase (the brightest available cue was always 16% sucrose, and the dimmest was always 1% sucrose).

We found that the preference for the higher-value cue decreased after a luminance shift, suggesting that changes in visual salience significantly impact value-based decision-making by rats. Results were separated by shift direction (upshift: 16 vs 4 LEDs; downshift: 4 vs 1 LED) and showed consistent reductions in choice preference and d’ (a measure of perceptual sensitivity derived from signal detection theory; Macmillan and Creelman, 2005) in both conditions. Using Drift Diffusion Models (DDMs), we found that luminance shifts reduced the drift rate (the speed of evidence accumulation) without affecting the decision threshold. These results clarify our previous findings (White et al., 2024; Palmer et al., 2024) and demonstrate that relative salience is a primary determinant of the efficiency of sensory evidence accumulation. Crucially, the computational effects of the luminance shifts show a dissociation from findings in Palmer et al. (2024), where prefrontal cortex inactivation affected the decision threshold but not the drift rate. Together, the results of the present study and Palmer et al. (2024) provide evidence for a mechanistic distinction, suggesting that while the prefrontal cortex controls the caution of the decision (threshold), visual salience controls the speed of sensory processing (drift rate).

## Materials & Methods

Procedures were approved by the Animal Care and Use Committee at American University (Washington, DC) and conformed to the standards of the National Institutes of Health Guide for the Care and Use of Laboratory Animals.

### Animals

Nine female Long Evans rats (200-250g, Charles-River) and five male Long Evans rats (300-400g, Charles River) were used in this study. Animals were individually housed on a 12/12 h light/dark cycle. During training and testing, animals had regulated access to food (12-16 grams) to maintain body weights at ∼90% of their free-access weights.

### Behavioral apparatus

Animals were trained in sound-attenuating behavioral boxes (Med Associates) that had a single, horizontally placed spout mounted to a lickometer 6.5 cm from the floor with a single white LED placed 4 cm above the spout. Solution lines were connected to 60cc syringes and solution was made available to animals by lick-triggered, single speed pumps (PHM-100; Med Associates) which drove syringe plungers. Each lick activated a pump which delivered roughly 30 μL per 0.5 second activation. On the wall opposite the spout, three 3D-printed nosepoke ports were aligned 5 cm from the floor and 4 cm apart, and contained Adafruit 3-mm IR Break Beam sensors. A Pure Green 1.2” 8×8 LED matrix (Adafruit) was placed 2.5 cm above the center of each of the three nose poke ports outside of the box for visual stimulus presentation. LED matrices are controlled using Adafruit_GFX and Adafruit_LEDBackpack libraries, and microcontroller software is provided in a previous publication (Swanson et al., 2021).

### Training procedure: Cue-Value Learning

Animals were trained in a behavioral task previously described (White et al. 2024; Palmer et al. 2024) (Figure 1A). First, animals licked at a reward spout in an operant chamber to receive 16% wt/vol liquid sucrose with the LED above the spout turned on. In subsequent sessions, animals were trained using the method of successive approximations to respond in nose poke ports in response to distinct visual stimuli. A 4×4 square of illuminated LEDs displayed over the center port signaled animals to initiate trials. Trial initiation was followed by lateralized presentation of one of two cues, which will be referred to as “single-offer” trials (Figure 1B, top). The cues are referred to as the high-luminance (16 illuminated LEDs) or low-luminance (one illuminated LED) throughout this paper. They were presented randomly by side (left or right). The location of illuminated LEDs changed every millisecond during the period of illumination, which began when rats entered the central port and ended when rats entered one of the lateralized ports. Responses for high-luminance yielded access to 16% wt/vol sucrose reward at the reward spout, and low-luminance yielded a 1% wt/volume sucrose reward. Responses to non-illuminated ports were considered as errors and were unrewarded. Responses that took longer than 5 seconds following trial initiation were counted as errors of omission and were unrewarded. On valid trials, animals had to collect their reward within 5 seconds following responses to receive fluid. Animals were trained for five, 60-minute sessions of at least 100 trials in single-offer sessions.

**Figure 1.**
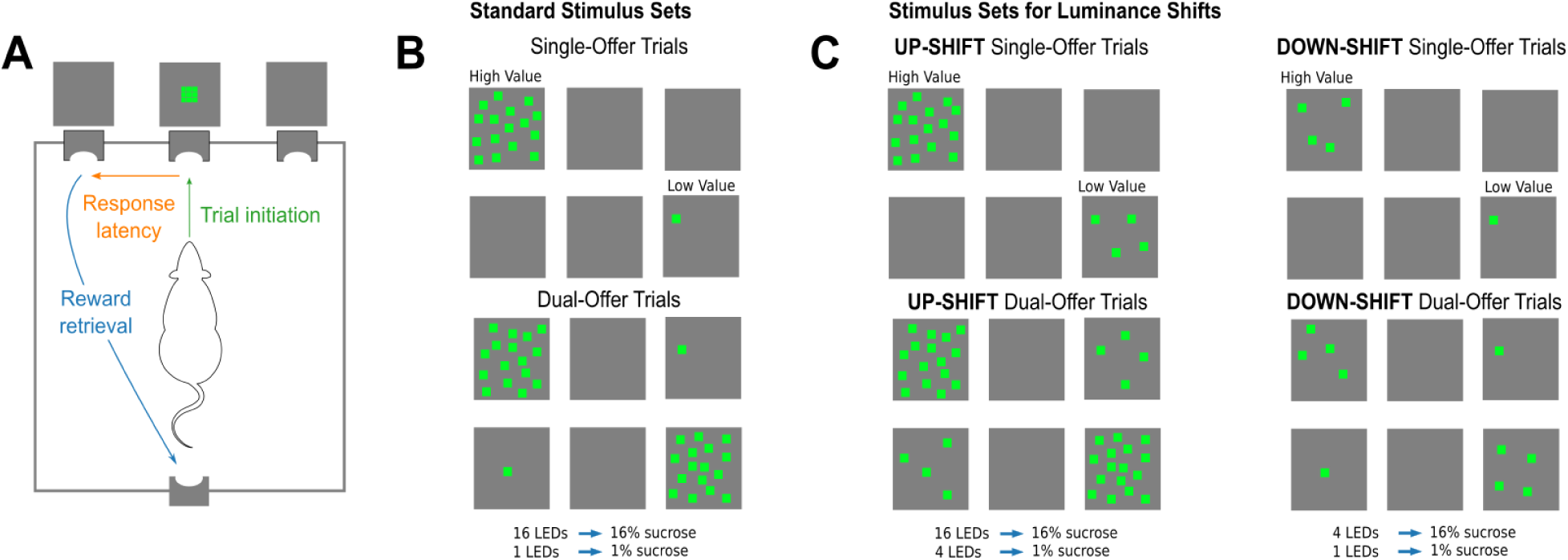
Experimental design. The behavioral apparatus and the visual stimulus pairings are shown. The visual stimuli were used to examine how graded changes in relative salience affect decision dynamics within a fixed reward structure, using a two-alternative forced-choice (2AFC) task. **A.** Illustration of the operant chamber and behavioral sequence comprising a trial in the task. Rats initiated trials by entering a central port. Stimuli were presented on LED grids located immediately above two lateralized choice ports. Rats entered one of the choice ports to select a stimulus. Then, they traveled to a reward port, located on the opposite wall of the testing chamber, to collect a liquid sucrose reward. **B.** Standard Stimulus Sets: Stimuli as they appear on LED grids during cue-value training and the control sessions reported in this study. Responding to 16 LEDs yields 16% wt/vol sucrose, and 1 LED yields 1% wt/vol sucrose. **C.** Stimulus Sets with Luminance Shifts: Left: Stimulus set for up-shift testing. Responding to 16 LEDs yields 16% wt/vol sucrose, and 4 LEDs yields 1% wt/vol sucrose. Right: Stimulus set for down-shift testing. Responding to 4 LEDs yields 16% wt/vol sucrose, and 1 LED yields 1% wt/vol sucrose. Reward values were held constant across all shift conditions.

Following single-offer training, animals then underwent dual-offer training. In 60-minute sessions, two-thirds (∼67%) of trials were single-offer trials, as described above. In the remaining third (∼33% of trials), animals were presented with both the high- and low-luminance cues simultaneously, randomized by side, and the animals had the choice of responding to either cue. These trials are referred to as “dual-offer” trials (Figure 1B, bottom). Single- and dual-offer trials were interleaved throughout the 60-minute sessions. Animals experienced five 60-minute dual offer sessions before undergoing behavioral testing. Training procedures in the present study are consistent with that described in Palmer et al. (2024). A difference from White et al. (2024) is that the animals only experienced choice sessions during this stage of training. In the study by White et al. (2024), there were sessions with only single offer trials interleaved between sessions with choice learning. All test sessions following training included mixtures of single-offer and dual-offer trials, with single-offer and dual-offer trials each comprising approximately 50% of trials. This proportion was increased relative to training to collect more dual-offer choice data during testing.

### Behavioral Testing: Between-Session Cue Shifting

In previous studies from our group (White et al. 2024; Palmer et al. 2024), cue luminance was congruent with reward value, where the brighter cue yields a higher concentration of liquid sucrose. This makes it difficult to determine how much each factor independently contributes to choice behavior. To examine this, we modified the task to introduce a new cue with an intermediate luminance, testing how changes in relative salience affect decision dynamics within a fixed reward structure. This cue will be referred to as the mid-luminance cue (4 illuminated LEDs).

Following the fifth dual-offer training session, animals were run in one additional session with high and low luminance cues to serve as a behavioral control. The following day, animals were run in a session where one cue was switched out and replaced with the mid-luminance cue. The reward value of this stimulus was the same as the one removed from the procedure. An “up-shift” in luminance refers to sessions where the low luminance cue is replaced, so rats will choose between the high- and mid-luminance cue. During up-shifts, responding to the high-luminance cue yields 16% sucrose, and the mid-luminance yields 1% sucrose. A “down-shift” in luminance refers to sessions where the high-luminance is replaced, so rats will choose between the mid- and low-luminance cues. During down-shifts, the mid-luminance cue yields 16% sucrose, and the low-luminance cue yields 1% sucrose. Up-shifts and down-shifts are shown in Figure 1C. All animals experienced two up-shift and two down-shift sessions. Each shift session was interleaved with a control session of the original high and low luminance cues.

The behavioral measures of interest are referred to as “latency,” “choice preference,” “detection,” “reward retrieval,” and “inter-trial interval.” Latency is measured as the time taken for rats to go from trial initiation to responding in an illuminated choice port after the onset of the visual stimuli. Only valid trials with latencies that were less than five seconds were included in analyses. Choice preference is measured as the ratio of dual-offer trials in which the animals responded to the higher luminance cue (high luminance cue in control and up-shifts, and mid luminance cue in down-shifts). Detection is the ratio of correct responses in single-offer trials. Reward retrieval is measured as the time it takes from making a nose poke response to the time that reward is collected. Inter-trial interval (ITI) is measured as the time from one trial initiation until the next trial initiation.

### Behavioral Testing: Within-Session Cue Shifting

Following between-session cue shifts, animals then underwent within-session cue shifts. In these sessions, animals would complete 100 trials with high and low luminance cues, and then either an up- or down-shift would occur in the middle of the session and would persist for the remainder of the 60-minute session. Animals experienced two within-session up-shifts and two within-session down-shifts. Each within-session shift was interleaved with a control session of the original high and low luminance cues. Behavioral measures of interest are the same as those listed above. One female rat did not perform enough post-shift trials (<10) to be included in analyses. The remaining 13 rats (eight females, five males) were included in within-session cue shifts.

Because animals could perform any number of trials-post shift, we wanted to determine whether any behavioral effects were impacted by the number of post-shift trials completed. We looked at effects in only animals who completed 100 or more post-shift trials, effects only up to the first 100 post-shift trials (including animals who did not reach 100 post-shift trials), and effects of animals who completed 100 or more post-shift trials, but only the first 100 trials. We found no differences in effects based on the number of post-shift trials, so all reported effects include all post-shift trials that each animal completed. The behavioral training and testing timeline is summarized in Figure 1.

### Statistical Analyses

Behavioral events were recorded through MED-PC and extracted through custom scripts written in Python (Anaconda distribution: https://www.continuum.io/). Statistical analyses were carried out using the pingouin package for Python (Vallat, 2018) and scipy (Virtanen et al., 2020). Repeated measures ANOVAs were used to assess effects on choice preference, latency, and d’ (Macmillan and Creelman, 2005). Shift direction (upshift vs downshift) was analyzed separately throughout; results were not combined across directions. Within-subject effects included session (control versus shift) and, for within-session analyses, period (pre versus post shift). Latencies were logarithmically transformed to account for skewed distributions. Reported statistics include F-statistics and p-values. Greenhouse-Geisser corrections were applied where sphericity was violated. Where applicable, pairwise post hoc comparisons were performed with Holm corrections (pingouin: pairwise_tests with padjust=“holm”).

We conducted an analysis of asymmetry between latencies as follows: the brighter-minus-dimmer latency difference was computed per animal per session and submitted to RM-ANOVA with session as the within-subject factor. To evaluate perceptual performance, d’ (a bias-free measure of perceptual sensitivity) was computed as sqrt(2) * z(p_correct), where p_correct is the proportion of dual-offer trials on which the animal chose the higher-value cue, clipped to [0.001, 0.999] to avoid infinite values. We assessed the stability of shift effects across repeated exposures using paired t-tests (scipy: ttest_rel) comparing each animal’s first and second exposure to each shift direction. For all statistical analyses, each rat was dropped from tests to confirm that effects were observed across animals.

To assess whether shift effects on choice preference were sustained across sessions, we compared performance in early (first 50 dual-offer trials) and late (last 50 dual-offer trials) blocks within each session type. A RM-ANOVA with session and block as within-subject factors was used, and pairwise comparisons of late-block preference across session types were Holm-corrected.

### Drift Diffusion Models

The HSSM package (version 0.3.0; https://doi.org/10.5281/zenodo.17247695) was used to quantify effects of session (between-session shifts: control vs shift) or period (within-session shifts: control post vs shift post) on decision making. We estimated four key parameters of drift diffusion models (DDMs): the drift rate, decision threshold, starting point bias, and non-decision time. Drift rate accounts for how quickly the rats integrate information about the stimuli. Threshold accounts for how much information is needed to trigger a decision. Starting point bias accounts for variability in the starting point of evidence accumulation. Non-decision time accounts for the time taken to initiate stimulus processing and execute the motor response (choice).

HSSM models were fit hierarchically and allowed for a single DDM parameter (drift rate, threshold, starting point bias, non-decision time) to vary freely over sessions. The other parameters were estimated globally. Models were fit to dual-offer trials only, as these represent genuine two-alternative choices. Two animals with atypically long response latencies (PB04 and PB05) were excluded because their RT distributions caused sampling instability; all DDM results are reported for n=12 animals. Models were fit separately for each shift direction (upshift and downshift) using the formula style “v ∼ 0 + session + (1|participant_id)”, which returns direct posterior estimates for each session level. The pymc sampler was used for all parameters. Models were fit by running version 0.3.0 of the package under Python 3.12. Models were run with 4 chains each, with 2000 tune and 2000 draw iterations. Target acceptance was increased to 0.90 from the default of 0.8 to improve convergence.

Convergence was validated based on the Gelman-Rubin statistic (Gelman and Rubin, 1992), which was below 1.01 for all parameters reported in this paper, with posterior predictive checks that compared response times from the rats and the models, and by plotting the probability density functions for the observed and predicted response times for each rat. The autocorrelations and distributions of the parameters and predictions of the response latency distributions for each animal were visually assessed to further assess convergence.

Statistics reported include the mean and 94% highest density interval (HDI: 3% and 97%) of the posterior distributions. Instances where the 94% HDIs associated with a given level of the factor (session) did not overlap with the mean parameter estimate of another level of the factor were considered as significantly different. Since HSSM uses regression-style parameter specification, we used custom Python scripts to extract fits for the base categorical level.

To validate the main findings of HSSM, we ran equivalent models using HDDM (Wiecki et al., 2013; version 0.9.6, Python 3.7) on the original collapsed dataset. This comparison is reported in Extended Data. HDDM models were fit with 50,000 samples, discarding the first 25,000 as burn-in and retaining every 10th sample (thinning of 10). Convergence was assessed using the Gelman-Rubin statistic and visual inspection of chain traces. Statistics from HDDM are reported as 95% credible intervals.

## Results

### Between-Session Shifts of Cue Luminance

The first experiment examined whether performance was driven by cues or values by replacing one of the trained cues with a novel cue of intermediate luminance while keeping rewards fixed. We tested rats with this cue in full behavioral sessions to determine how they would respond to the novel cue consistently over a session, without the influence of surprise that may come from introducing novel cues for the first time within-session. The rats underwent four rounds of between-session shifts, with two up-shifts and two down-shifts in an alternating pattern. Each round consisted of a control session (“Control 1”), a shift session with the intermediate cue, and a subsequent control session (“Control 2”). Results are reported separately for upshift (16 vs 4 LEDs) and downshift (4 vs 1 LED) conditions, averaged across the four rounds of testing.

Choice preference was significantly reduced during shift sessions relative to surrounding control sessions in both upshift and downshift conditions (Figure 2A). RM-ANOVAs with session as the within-subject factor revealed significant effects in both directions (Upshift: F(2,26)=64.9, p<0.001; Downshift: F(2,26)=46.4, p<0.001). Pairwise comparisons confirmed that preference was reduced during shift sessions relative to both Control 1 and Control 2 in both directions (all p<0.001, Holm-corrected). In upshift sessions, preference dropped from 83.2% in control to 66.5% during the shift. In downshift sessions, preference dropped from 80.0% to 71.1%. Preference remained well above chance in both conditions, consistent with value maintaining behavior above chance while relative salience determines the magnitude of the effect.

**Figure 2.**
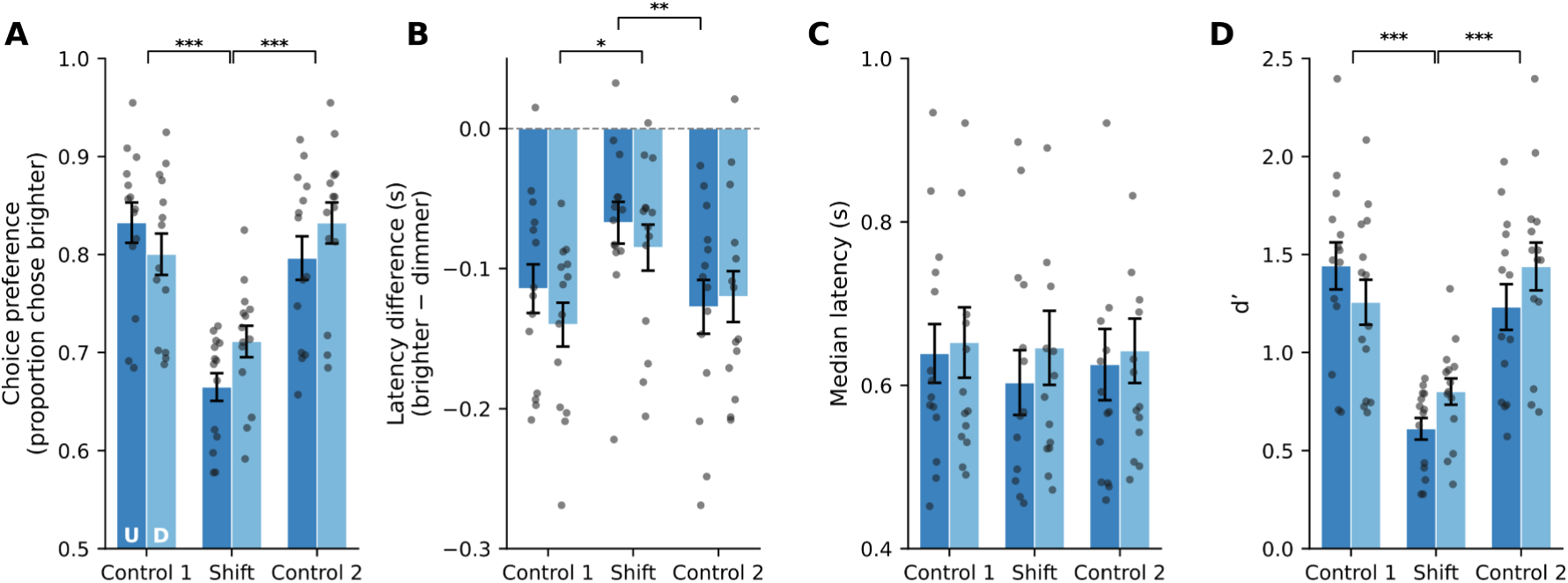
Behavioral effects of between-session luminance shifts. The mid-luminance cue (4 LEDs) replaced the low-luminance cue (1-LEDs) in upshift sessions (dark blue). In downshift sessions (light blue), the high-luminance cue (16 LEDs) was replaced with the 4-LED cue. Reward values were the same throughout. **A.** Choice preference was significantly reduced during shift sessions relative to both surrounding control sessions for both up and down shifts (all p<0.001, Holm-corrected). **B.** The brighter-minus-dimmer latency difference was reduced during shift sessions (Upshift: F(2,26)=5.90, p=0.008; Downshift: F(2,26)=4.14, p=0.028). Pairwise comparisons indicated a reduction in downshift sessions relative to Control 1 (p=0.040) and a delayed reduction in upshift sessions relative to Control 2 (p=0.009). **C.** Overall median latency did not differ significantly across session types for both types of shifts. This finding suggests that the session effect was specific to the brighter-dimmer latency difference rather than an overall change in response speed. **D.** d’ was significantly reduced during shift sessions relative to surrounding controls for both up and down shifts (all p<0.001, Holm-corrected). This finding suggests that graded changes in relative salience reduced the quality of the choice signal. Data shown here were averaged across four rounds of testing. Individual animal values are shown as dots. Error bars indicate SEM.

The brighter-minus-dimmer latency difference was reduced during shift sessions in both directions (Figure 2B). RM-ANOVAs confirmed significant session effects for upshift (F(2,26)=5.90, p=0.008) and downshift (F(2,26)=4.14, p=0.028) conditions. The timing of the effect differed by direction: in downshift sessions the latency gap was significantly reduced relative to Control 1 (p=0.040, Holm-corrected), whereas in upshift sessions the reduction was significant relative to Control 2 (p=0.009). This asymmetry suggests that replacing the dominant high-value cue with a less salient alternative disrupts latency immediately, while making the low-value cue more similar to the high-value cue produces a delayed carry-over effect. Overall median latency did not differ significantly across session types in either direction (Figure 2C), confirming that the session effect operated through the brighter-dimmer latency difference rather than a general change in response speed. There were no effects of the shifts on reward retrieval or ITI (not shown).

To quantify how cue similarity influenced the quality of the choice signal, we computed a d’ from dual-offer trial data (Figure 2D). d’ was significantly reduced during shift sessions relative to surrounding controls in both directions (all p<0.001, Holm-corrected), dropping from 1.44 to 0.61 in upshift sessions and from 1.26 to 0.80 in downshift sessions. These reductions are consistent with the decrease in drift rate observed in the DDM analyses below.

Single-offer trial accuracy was high across all session types and was unaffected by the luminance shifts (Figure 2-1A-B). Detection rates for both the brighter and dimmer cues remained near the ceiling in control and shift sessions alike, with no significant effect of session on detection for either cue in either shift direction (all p>0.05). Median latencies on single-offer trials were similarly unaffected by session (all p>0.05), though animals responded faster to the brighter cue regardless of session type, consistent with the luminance-driven latency difference observed in dual-offer trials. These results confirm that animals could identify each cue individually regardless of the luminance pairing in effect during that session. The shift effects on dual-offer choice preference and latency therefore reflect changes in the relative comparison between simultaneously presented options rather than a failure to detect either stimulus in isolation.

Drift diffusion models fit separately for each shift direction revealed a consistent pattern across conditions (Figure 3). Models were fit to dual-offer trials from the first shift exposure in each direction, excluding two animals with atypically long response latencies (PB04 and PB05; n=12). A comparison of HSSM and HDDM parameter estimates is provided in Figure 3-1.

**Figure 3.**
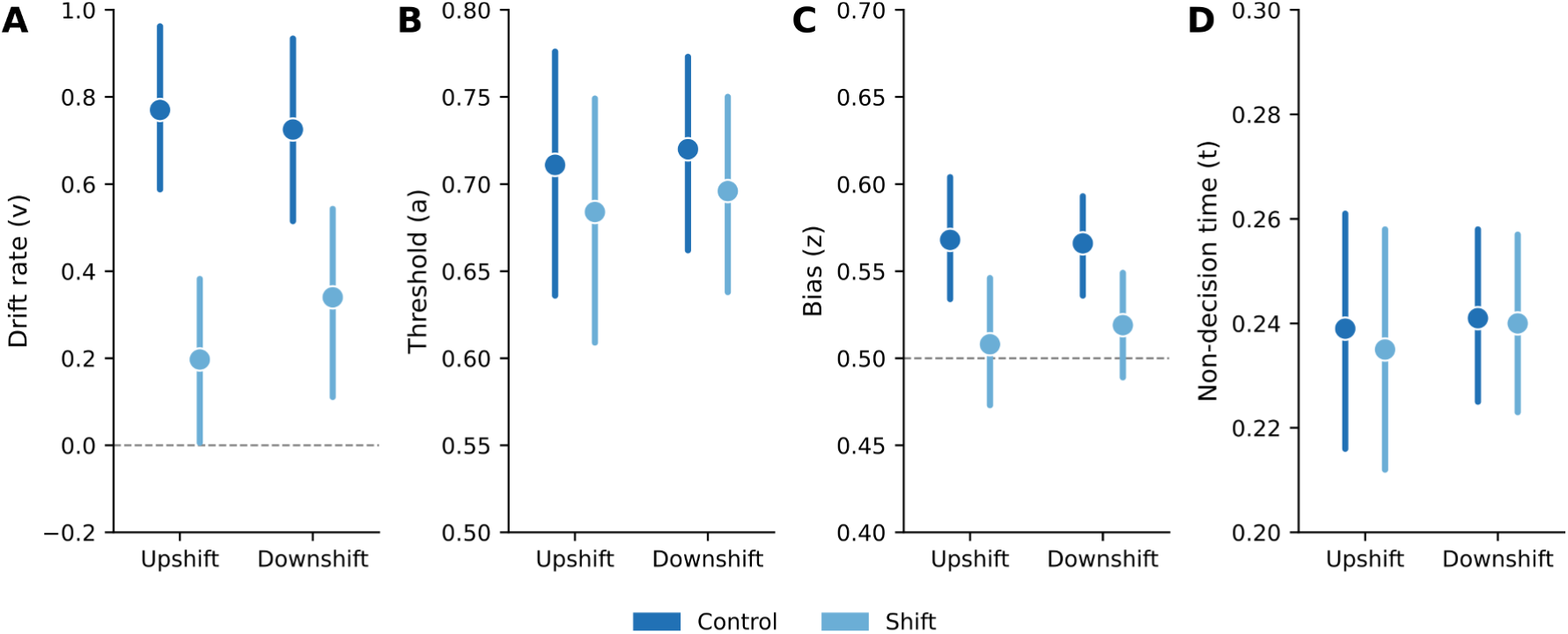
Drift diffusion modeling of the between-session luminance shifts. Parameters were estimated separately for upshift (16 vs 4 LEDs) and downshift (4 vs 1 LED) conditions using hierarchical drift diffusion models fit in HSSM (v0.3.0). Points show posterior means and lines show 94% highest density intervals (HDIs) for control (dark blue) and shift (light blue) sessions. Non-overlapping HDIs indicate a significant difference between conditions. **A.** Drift rate was reduced during shift sessions in both upshift and downshift conditions, indicating that reduced cue discriminability slowed the accumulation of sensory evidence. **B.** Decision threshold was unaffected by the luminance shifts in both conditions, consistent with animals maintaining the same level of decision caution across session types. **C.** Starting point bias was reduced during shift sessions, reflecting attenuation of the prior expectation toward the brighter, higher-value option when the two cues became more perceptually similar. The dashed line at 0.5 indicates no bias. **D.** Non-decision time was unaffected by the luminance shifts. Models were fit to dual-offer trials from the first shift exposure in each direction, excluding two animals with atypically long response latencies (PB04 and PB05; n=12).

Drift rate was significantly reduced during shift sessions in both upshift and downshift conditions, with non-overlapping HDIs between control and shift. For the downshift first exposure, control drift rate was 0.725 (HDI 0.515–0.934) and shift drift rate was 0.340 (HDI 0.111–0.543). Starting point bias was also reduced: control 0.566 (HDI 0.536–0.593) versus shift 0.519 (HDI 0.489– 0.549). Decision threshold was unaffected, with substantially overlapping HDIs: control 0.720 (HDI 0.662–0.773) versus shift 0.696 (HDI 0.638–0.750). Non-decision time was similarly stable: control 0.241s (HDI 0.225–0.258) versus shift 0.240s (HDI 0.223–0.257).

For the upshift first exposure, control drift rate was 0.77 (HDI 0.588–0.962) and shift drift rate was 0.197 (HDI 0.006–0.382). Starting point bias was also reduced: control 0.568 (HDI 0.534–0.604) versus shift 0.508 (HDI 0.473–0.546). Decision threshold was unaffected, with substantially overlapping HDIs: control 0.711 (HDI 0.636–0.776) versus shift 0.684 (HDI 0.609–0.749). Non-decision time was similarly stable: control 0.239s (HDI 0.216–0.261) versus shift 0.235s (HDI 0.212–0.258).

To assess whether the shift effect was sustained across the session, we compared choice preference in early (first 50 dual-offer trials) and late (last 50 dual-offer trials) blocks within each session type (Figure 4). RM-ANOVAs with session and block as within-subject factors revealed significant session effects in both directions (Upshift: F(2,26)=66.5, p<0.001; Downshift: F(2,26)=42.1, p<0.001) with no effect of block and no session × block interaction (all p>0.11). Late-block pairwise comparisons confirmed significant reductions during shift sessions relative to both control periods in both directions (all p<0.001, Holm-corrected). Early and late blocks did not differ within shift sessions (upshift: p=0.79; downshift: p=0.96), indicating that the shift effect was present from the first trials and did not adapt. A small but significant difference between the two surrounding control sessions was observed in both directions (upshift: p=0.037; downshift: p=0.008), likely reflecting session order effects across the four rounds of testing.

**Figure 4.**
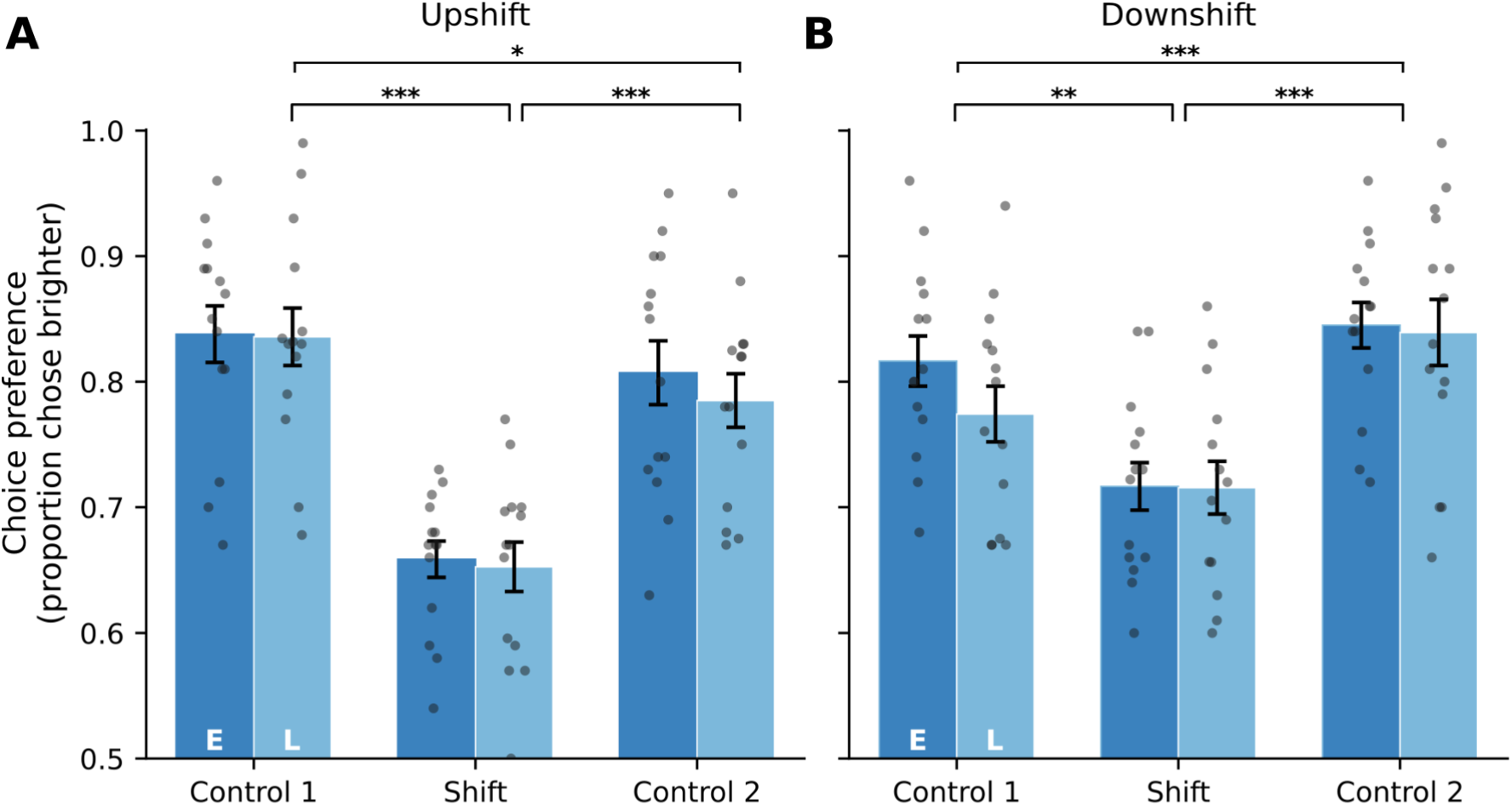
Split-session analysis of choice preference during between-session luminance shifts. Early and late blocks represent the first and last 50 dual-offer trials within each session. **A.** Upshift sessions (low-value cue increased from 1 to 4 LEDs). A repeated-measures ANOVA revealed a significant effect of session (F(2,26)=66.5, p<0.001) with no effect of block and no session × block interaction (both p>0.34). Late-block pairwise comparisons confirmed significant reductions in shift sessions relative to both control periods (both p<0.001, Holm-corrected), with a residual difference between control sessions (p=0.037). **B.** Downshift sessions (high-value cue reduced from 16 to 4 LEDs). The session effect was again significant (F(2,26)=42.1, p<0.001) with no block effect or interaction (both p>0.11). Late-block comparisons showed significant reductions in shift sessions relative to both control periods (both p<0.001, Holm-corrected) and a difference between control sessions (p=0.008). In both directions, early and late blocks did not differ within shift sessions (upshift: p=0.79; downshift: p=0.96), indicating that the shift effect was present from the first trials and did not adapt across the session. Individual animal values are shown as dots. Error bars indicate SEM.

### Within-Session Shifts of Cue Luminance

Having found evidence of the intermediate cue’s effects on performance, we next examined how inserting the cue within-session would affect performance. All animals experienced four within-session shifts, alternating between two up-shifts and two down-shifts. Each round consisted of a control session (“Control 1”), a shift session, and a subsequent control session (“Control 2”). Results are reported separately for upshift and downshift conditions, averaged across rounds. In shift sessions, the cues initially had high (16 LEDs) and low (1 LED) luminance, and after 100 trials, one cue was replaced with the mid-luminance cue (4 LEDs). In control sessions, the cues remained at high and low luminance throughout, with the “shift” still defined as the first 100 trials (“pre-shift”) and all trials after (“post-shift”).

Choice preference was significantly reduced during shift sessions relative to surrounding control sessions in both directions (Figure 5A; all p<0.001, Holm-corrected), replicating the between-session finding. The brighter-minus-dimmer latency difference narrowed during shift sessions in both directions (Figure 5B). Holm-corrected pairwise tests revealed significant differences between Control 1 and Shift sessions and between Control 1 and Control 2 sessions in both directions (all p<0.001), with an additional difference between Shift and Control 2 sessions (p<0.01). Overall median latency did not differ significantly across session types in either direction (Figure 5C). d’ was significantly reduced during shift sessions relative to both surrounding control sessions in both directions (Figure 5D; all p<0.001, Holm-corrected).

**Figure 5.**
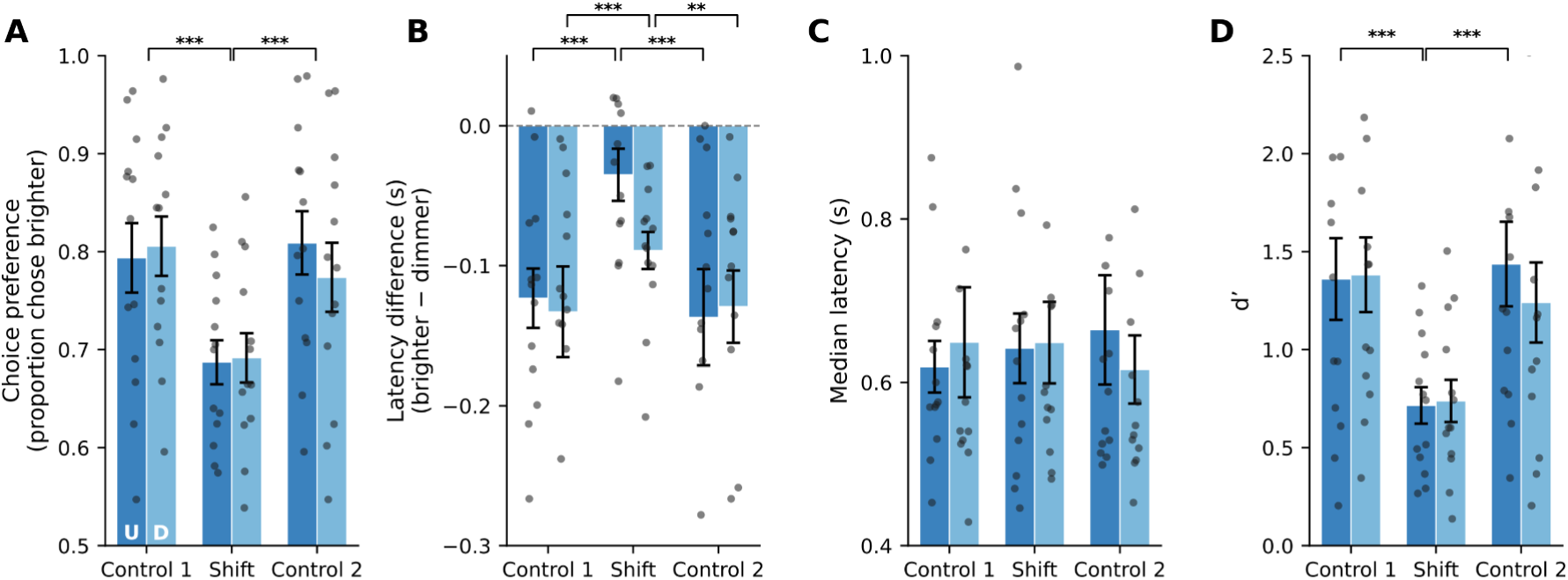
Behavioral effects of within-session luminance shifts. Upshift (dark blue) and downshift (light blue) conditions are shown side by side for each session type. **A.** Choice preference was significantly reduced during shift sessions relative to both surrounding control sessions in both directions (all p < 0.001, Holm-corrected). **B.** The brighter-minus-dimmer latency difference narrowed during shift sessions. Holm-corrected pairwise tests revealed significant differences between Control 1 and Shift sessions and between Control 1 and Control 2 sessions in both directions (all p < 0.001), with an additional difference between Shift and Control 2 sessions (p < 0.01). **C.** Overall median latency did not differ significantly across session types in either direction. **D.** d’ was significantly reduced during shift sessions relative to both surrounding control sessions in both directions (all p < 0.001, Holm-corrected). Individual animal values are shown as dots. Error bars indicate SEM.

As in the between-session experiment, single-offer detection rates and latencies were unaffected by within-session luminance shifts in either direction (Figure 2-1C-D; all session effects p>0.05). These findings confirm that the post-shift reduction in dual-offer choice preference reflects a change in the comparison between simultaneously presented cues rather than impaired detection of individual stimuli.

Drift diffusion models were fit to post-period dual-offer trials, comparing the post-shift period in shift sessions to the equivalent post-period in control sessions (Figure 6). The first 100 post-period trials per animal were used to match the fixed 100-trial pre-shift window. Models were fit to n=12 animals (excluding PB04 and PB05).

**Figure 6.**
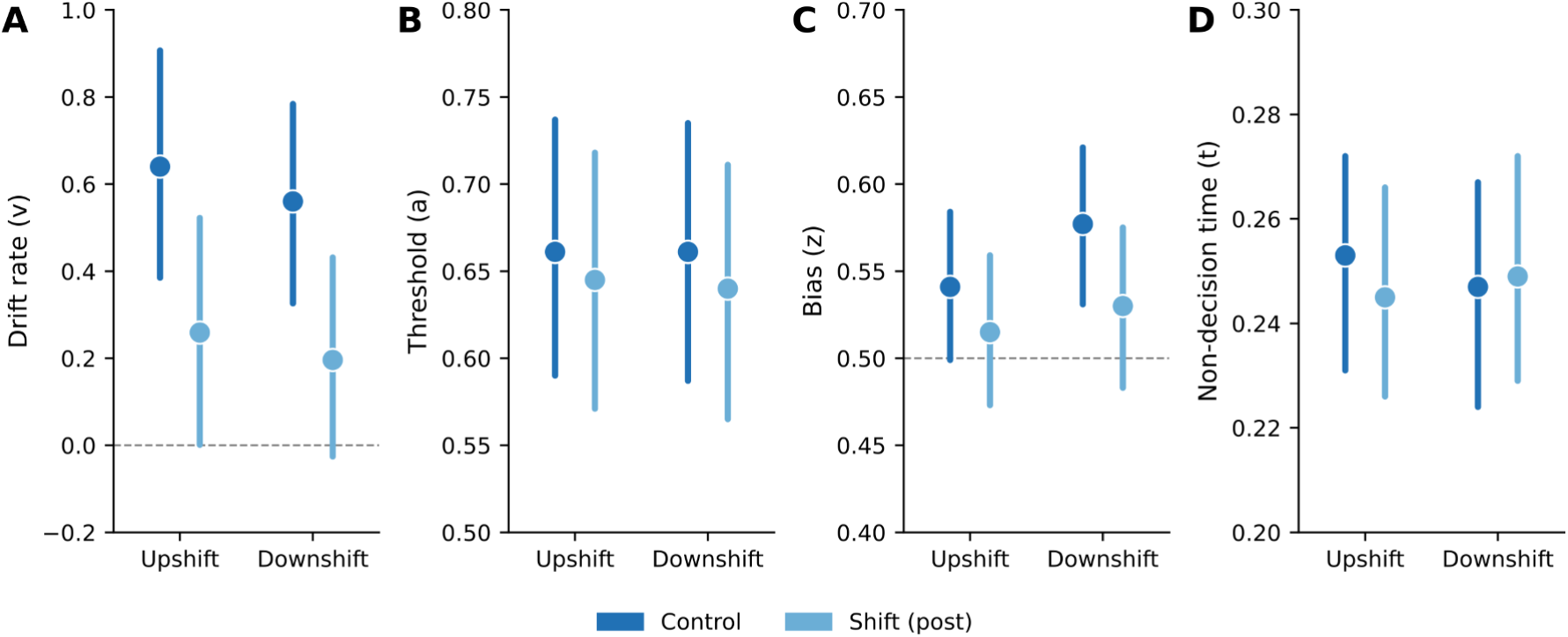
Drift diffusion model parameter estimates for within-session luminance shifts. Parameters were estimated from post-period trials in shift sessions compared to the equivalent post-period trials in control sessions, isolating the effect of the luminance change from general within-session factors. **A.** Drift rate was reduced in the post-shift period relative to the equivalent control period in both upshift and downshift conditions. In the downshift condition, the lower bound of the shift HDI approached zero, indicating greater variability in the magnitude of the drift rate reduction across animals. **B.** Decision threshold was unaffected by the within-session luminance shifts. **C.** Starting point bias was reduced in the post-shift period in both conditions, with shift HDIs approaching the 0.5 no-bias reference line. **D.** Non-decision time was stable across conditions. Models were fit to the first 100 post-period dual-offer trials per animal to match the fixed 100-trial pre-shift window and balance trial counts across conditions (n=12).

Drift rate was reduced in the post-shift period relative to the equivalent control period in both directions, with non-overlapping HDIs (downshift control: 0.56; downshift shift: 0.196; upshift control: 0.64; upshift shift: 0.259). In the downshift condition, the lower bound of the shift HDI approached zero, indicating greater variability in the magnitude of the drift rate reduction across animals. Decision threshold was unaffected in both directions. Starting point bias was reduced in the post-shift period in both conditions (downshift control: 0.577; downshift shift: 0.53; upshift control: 0.541; upshift shift: 0.515), with shift HDIs straddling the 0.5 no-bias reference line. Non-decision time was stable across conditions.

### Effects of Repeated Exposures to the Shift Procedures

To assess whether the shift effect changed with repeated exposure to the novel cue, we compared each animal’s choice preference between their first and second experience of each shift direction, across both experiments (Figure 7). For between-session shifts, choice preferences did not differ between exposures for either upshift (F(1,13)=0.54, p=0.48, η²g=0.010) or downshift (F(1,13)=0.19, p=0.67, η²g=0.003) conditions. The same pattern was observed for within-session shifts, with no exposure effect for upshift (F(1,12)=0.22, p=0.65, η²g=0.008) or downshift (F(1,12)=0.18, p=0.68, η²g=0.004). The shift effect was of comparable magnitude on the first and second exposure in all conditions, indicating that animals did not adapt to repeated luminance shifts across the four rounds of testing. This result argues against the possibility that animals were disregarding the novel cue or gradually acquiring a new cue-value discrimination.

**Figure 7.**
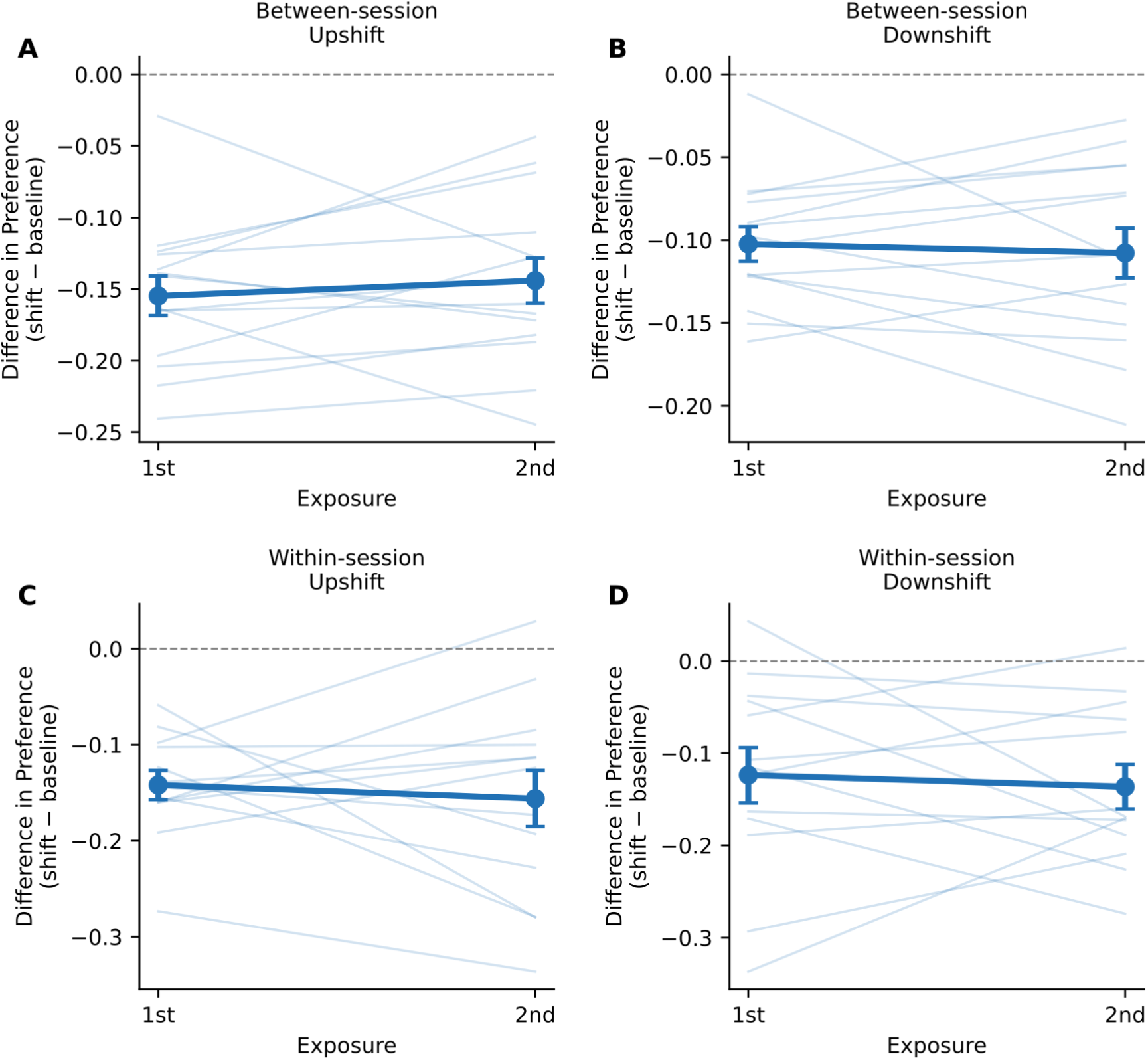
Stability of shift effects on choice preference across repeated exposures. Each panel shows the difference in choice preference during shift sessions relative to the mean of the surrounding control sessions (shift minus baseline) for each animal’s first and second exposure to each shift direction. **A.** Between-session upshift. **B.** Between-session downshift. **C.** Within-session upshift. **D.** Within-session downshift. The reduction in choice preference during shift sessions was consistent across first and second exposures in all four conditions (all p>0.46, paired t-tests), indicating that the effect of reduced cue discriminability on choice behavior did not diminish with repeated experience. This result argues against the possibility that animals disregarded the novel mid-luminance cue or gradually acquired a new cue-value discrimination across exposures. Individual animal trajectories are shown in light blue. Dark blue points show the group mean ± SEM. The dashed line at zero indicates no change from baseline.

To examine whether the direction of the shift influenced the latency effect across experiments, we computed the reduction in the brighter-minus-dimmer latency gap (shift minus control) for each animal’s first and second exposure to each direction (Figure 8). No significant differences between exposures were observed in any condition (all p>0.05), confirming that the latency effect was also stable across repeated exposures. Minor differences between upshift and downshift conditions were observed in the within-session experiment, with the upshift producing a somewhat larger latency delta than the downshift on the second exposure. These differences did not reach statistical significance and are consistent with the directional asymmetry in the brighter-minus-dimmer latency effect described for the between-session experiment above.

**Figure 8.**
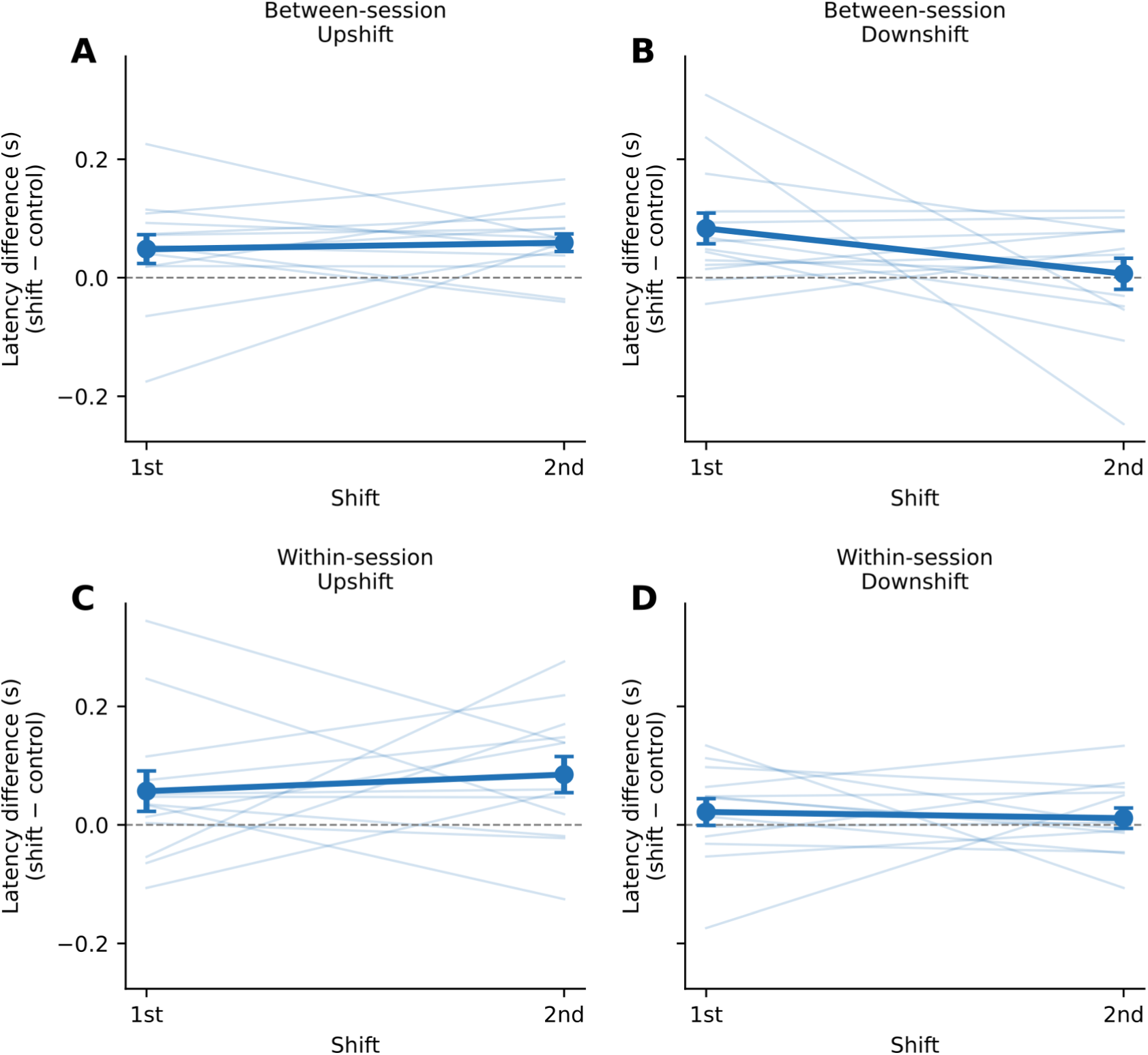
Latency asymmetry across repeated shift exposures. Each panel shows the difference in the brighter-minus-dimmer latency difference between shift and control sessions for the first and second exposure to each shift direction. **A.** Between-session upshift: latency delta was small and stable across exposures. **B.** Between-session downshift: latency delta was positive at the first exposure and decreased toward zero at the second. **C.** Within-session upshift: latency delta was stable across exposures. **D.** Within-session downshift: latency delta remained near zero across both exposures. No significant differences between exposures were observed in any condition (all p>0.05). Minor differences between upshift and downshift conditions in the within-session experiment are described in the Results. Individual animal data are shown in light blue. Dark blue points and lines show the group mean ± SEM. The dashed line at zero indicates no difference from control.

## Discussion

We investigated whether the brightness of cues or the rewards they signal is mainly responsible for the behavioral differences seen in previous studies of value-guided decisions by rats (White et al., 2024; Palmer et al., 2024). Using the same task design, we trained rats to respond to a bright cue (16 LEDs) for a high concentration of liquid sucrose (16% wt/vol) and to a dim cue (1 LED) for a low concentration (1% wt/vol). To examine effects of relative salience, we replaced one of the cues with a mid-luminance cue (4 LEDs) and assigned it the same reward value as the original cue. This manipulation tested how changes in relative salience affect decision dynamics within a fixed reward structure. This replacement resulted in either an upshift (mid-luminance replacing low-luminance, yielding 1% sucrose) or a downshift (mid-luminance replacing high-luminance, yielding 16% sucrose). Results were analyzed separately for each shift direction across both between-session and within-session experiments.

By making luminance more similar, we found that the rats’ behavioral preferences seen in baseline sessions were significantly reduced in both directions, evident in changes to choice preference, d’, and the brighter-minus-dimmer latency difference. These results suggest that the perceptual discriminability of the cues is a primary determinant of choice behavior. Using drift diffusion modeling, we found that the changes in performance were driven by a decrease in the rate of sensory evidence accumulation, demonstrating a direct computational link between a low-level visual property and the dynamics of decision-making.

### Shifts in Luminance Reduce Behavioral Differences Between the Cues

Replacing one cue with a cue of intermediate brightness (4 LEDs) diminished the behavioral differences between the relatively brighter and dimmer cues. Choice preference for the brighter cue decreased (Figure 2A) and the brighter-minus-dimmer latency difference was reduced (Figure 2B) in both shift directions. The direction of the shift modulated the temporal dynamics of the latency effect: in downshift sessions the gap narrowed immediately relative to Control 1, whereas in upshift sessions the reduction was apparent relative to Control 2, suggesting a carry-over effect. Overall median latency did not change across session types (Figure 2C). These effects were paralleled by reductions in d’ (Figure 2D), confirming that graded changes in relative salience reduced the quality of the choice signal. Although rats still responded faster and more consistently to the relatively brighter cue during the shifts, the magnitude of this effect decreased, revealing that the salience of the cue itself plays a critical role in the behavioral effects previously reported in White et al. (2024) and Palmer et al. (2024).

We used Drift Diffusion Modeling (DDM) to understand how shifts in luminance affected the decision process. Across both between-session and within-session shifts, we consistently found that the drift rate decreased following shifts in cue luminance (Figures 3 and 6). As the drift rate represents the speed of sensory evidence accumulation, the reduction in drift rate suggests that when the cues become more perceptually similar, the rate at which the visual information is processed slows down. This is consistent with DDM literature, which links a reduction in drift rate to increased difficulty or reduced discriminability between stimuli (Ratcliff and McKoon, 2008; Myers et al., 2022). Shifting from a high luminance difference (16 vs. 1 LED, high signal-to-noise) to a lower difference (16 vs. 4 or 4 vs. 1 LED, low signal-to-noise) inherently amplifies the internal perceptual noise experienced by the animal, leading to a shallower slope of evidence accumulation. This slower, noisier evidence accumulation likely accounts for the reduction in choice preference for the brighter, higher-value cue.

The luminance shifts were also accompanied by a reduction in the starting point bias (Figures 3 and 6). Since reward values remained fixed, the reduction in bias suggests that when the cues were perceptually more similar, the animals were less biased towards the relatively brighter option prior to evidence accumulation. Although bias is traditionally influenced by the magnitude of reward (Ratcliff and McKoon, 2008; Mulder et al., 2012), our results suggest that in the absence of a large perceptual difference, this value-driven bias is attenuated by shifts in cue luminance. The reduction in bias helps explain the diminished latency differences between cues during the shifts, as a starting point further from the boundary necessitates a larger volume of evidence for a decision (Myers et al., 2022). The joint reduction in drift rate and starting point bias across both shift directions and both experimental designs is consistent with a salience-driven account in which perceptual discriminability influences both the rate of evidence accumulation during each trial and the prior expectation with which accumulation begins.

### Shift Effects Are Immediate and Sustained Within Sessions

To determine whether the shift effects reflected sensitivity to the change in relative salience or gradual acquisition of a new cue-value discrimination, we examined choice preference in early and late blocks within each session type (Figure 4). The reduction in choice preference was present from the first trials of each shift session and did not diminish across the session, with no significant early-versus-late difference within shift sessions in either direction (upshift: p=0.79; downshift: p=0.96). This pattern argues against the possibility that animals required experience with the novel cue to adjust their behavior. Instead, the immediate and sustained nature of the effect is consistent with the interpretation that reduced cue discriminability acts directly on the evidence accumulation process rather than triggering a learning process.

### Shift Effects Are Stable Across Repeated Exposures

A key alternative interpretation is that animals disregarded the novel mid-luminance cue or gradually learned a new cue-value discrimination across the four rounds of testing. We addressed this by comparing each animal’s first and second exposure to each shift direction in both experiments (Figures 7 and 8). The reduction in choice preference was consistent across exposures in all conditions, with no significant change from first to second exposure in either direction or either experiment (all p>0.46). The magnitude of the latency effect was similarly stable across exposures (all p>0.05). Had animals been disregarding the novel cue, the shift effect would have been smaller on first exposure and absent on subsequent ones. Had animals been acquiring a new discrimination, the shift effect would have diminished as they learned the new cue-value contingency. Neither pattern was observed. The stability of the effect across exposures confirms that it reflects a direct and immediate consequence of reduced cue discriminability on the evidence accumulation process rather than a learning or adaptation phenomenon.

### Visual Salience and Prefrontal Control of Value-Guided Decisions: A Double Dissociation

The primary finding that shifts in luminance reduce performance by decreasing the drift rate provides a crucial computational distinction regarding the neural control of value-based decisions. This result contributes to the emerging concept of anatomical-computational mapping, suggesting that the decision process is governed by modular neural components.

In our related study (Palmer et al., 2024), we found that reversible inactivation of the prelimbic cortex, part of the rodent prefrontal cortex (Laubach et al., 2018), in rats performing the same task selectively affected the decision threshold, leading to more impulsive choices, but did not affect the drift rate. The current study demonstrates the inverse: manipulating the visual salience of the stimulus reduces drift rate and starting point bias without affecting the decision threshold. This pattern was consistent across upshift and downshift directions and across both between-session and within-session experiments. This double dissociation suggests that the decision process is governed by two functionally distinct mechanisms. The salience (or discriminability) of the stimulus determines how efficiently evidence is accumulated, likely controlled by input from the visual system. The prefrontal cortex determines the animal’s cautiousness or decision policy, exerting top-down executive control over the amount of evidence required to make a choice.

This distinction is vital for dissecting the neural circuits underlying the separate computational components of decision-making. If the prelimbic cortex controls the decision threshold, other brain regions must control the drift rate, possibly including the more caudal anterior cingulate cortex (ACC) (Vazquez et al., 2024). The rodent ACC integrates visual, reward, and motor information (Huda et al., 2020), making it a strong candidate for controlling the sensory processing that underlies drift rate. Future studies could reversibly inactivate the rostral prelimbic and caudal ACC in the same animals, determining whether these neighboring cortical regions indeed have distinct effects on the decision process, thus testing the proposed modular framework for value-guided decisions.

## Author Note

### Contributions

Jensen Palmer, Kevin Chavez Lopez, and Mark Laubach designed the experiments. Jensen Palmer and Kevin Chavez Lopez carried out the experiments. Jensen Palmer analyzed the behavioral data and performed the drift diffusion modeling. Jensen Palmer, Kevin Chavez Lopez, and Mark Laubach wrote the manuscript. Some computer code for data analysis and graphical summaries was developed with Claude (Anthropic) as AI coding assistant. Claude was also used to perform checks on grammar and spelling and the accuracy of reported statistics and figure citations in the text.

## Acknowledgments

We thank Yogita Chudasama, David Kearns, and Fany Messanvi for helpful comments on the manuscript.

## Extended Data

**Figure 2-1.**
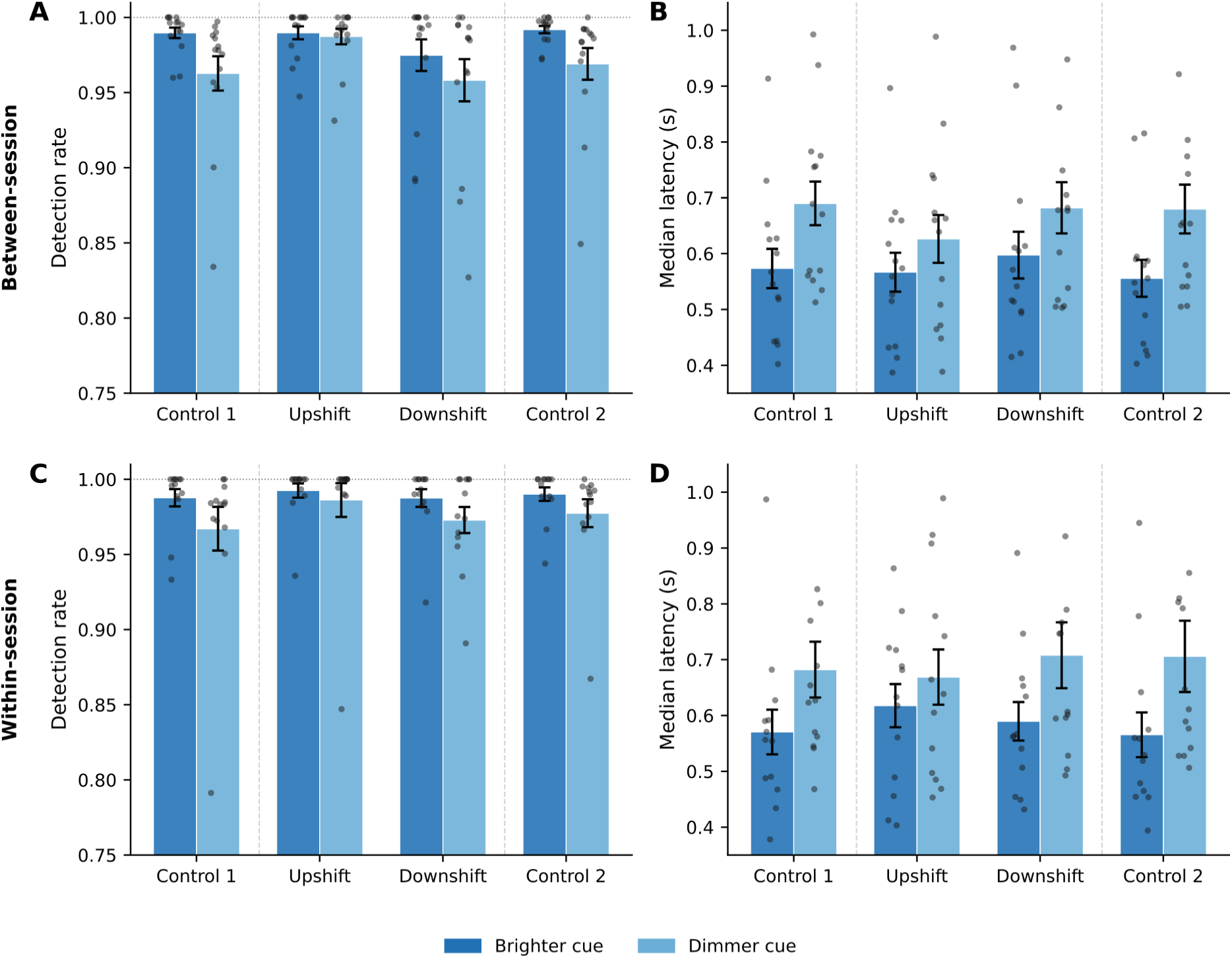
Single-offer trial performance was unaffected by luminance shifts. Detection rates and median latencies for single-offer trials are shown for control sessions (Control 1 and Control 2), upshift sessions (16 vs 4 LEDs), and downshift sessions (4 vs 1 LEDs). Brighter cue (dark blue) and dimmer cue (light blue) are shown as separate bars for each session type. **A.** Between-session detection rates. Detection remained near the ceiling across all session types for both cues, with no significant effect of session on detection in control sessions (p=0.061) and no significant difference between brighter and dimmer cues within shift sessions (both p>0.40). **B.** Between-session median latencies. No significant session effect was observed in control sessions (p=0.35). Animals responded faster to the brighter cue in upshift sessions (p=0.010), consistent with the luminance-driven latency advantage seen in dual-offer trials. **C.** Within-session detection rates (post-shift period). There was no significant effect in control sessions (p=0.088) and no significant brighter-versus-dimmer difference in upshift sessions (p=0.39). A significant brighter-versus-dimmer difference was present in downshift sessions (p=0.006), indicating that animals could detect the 4-LED versus 1-LED distinction. **D.** Within-session median latencies (post-shift period). No significant session effect was found in the control sessions (p=0.44). Significant brighter-versus-dimmer latency differences were present in downshift sessions (p=0.002). Individual animal values are shown as dots. Error bars indicate SEM.

**Figure 3-1.**
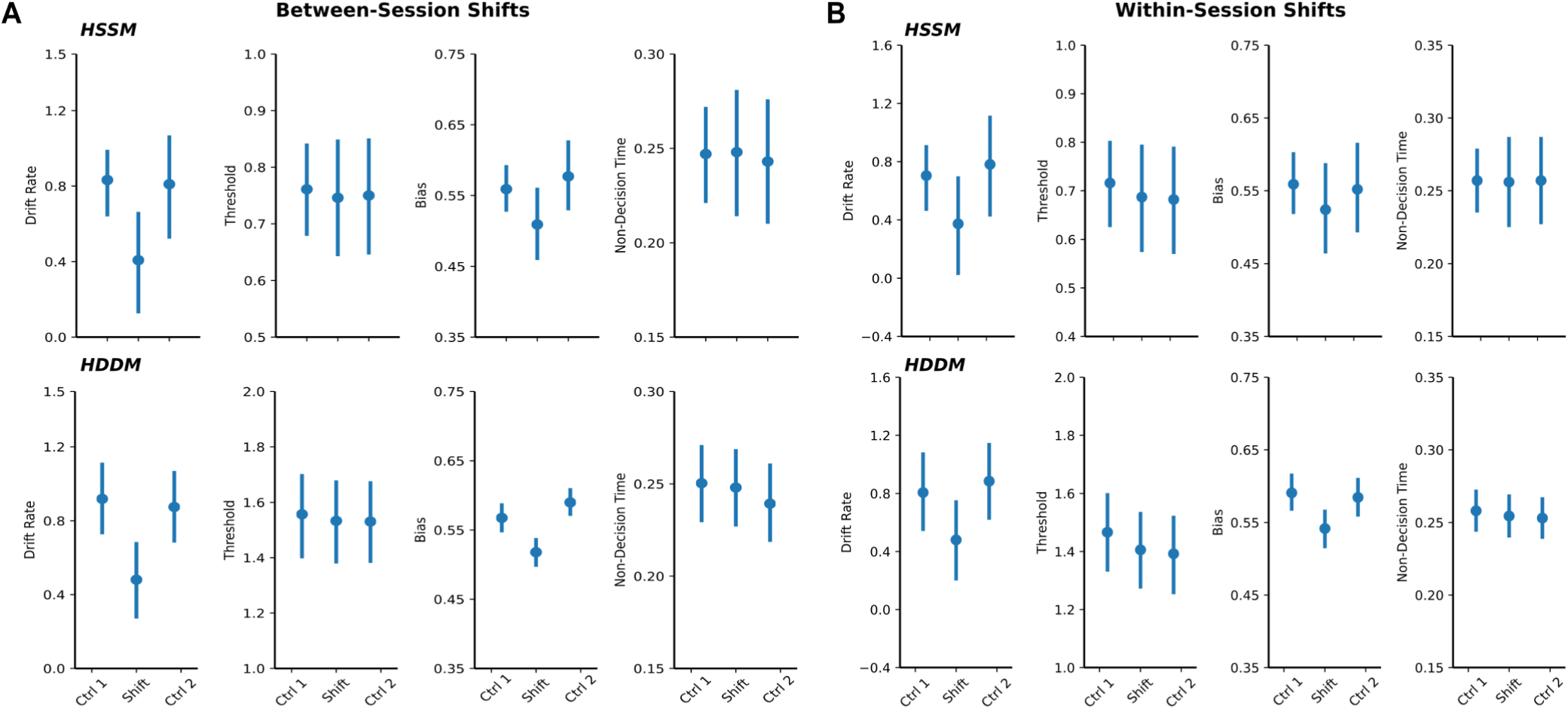
Comparison of HSSM and HDDM drift diffusion model parameter estimates. Results are shown for between-session shifts (A) and within-session shifts (B), collapsed across upshift and downshift directions. Each panel shows posterior means and highest density intervals (HDIs) for control sessions (Ctrl 1, Ctrl 2) and shift sessions across four DDM parameters: drift rate, decision threshold, starting point bias, and non-decision time. Top rows: HSSM (v0.3.0) estimates with 94% HDIs. Bottom rows: HDDM (v0.9.6) estimates with 95% credible intervals. Both packages show consistent reductions in drift rate and starting point bias during shift sessions, with threshold and non-decision time unaffected. Note that threshold estimates differ in absolute scale between packages due to differences in parameterization, but the pattern of effects is consistent.

